# Integrative taxonomy using traits and genomic data for Species Delimitation with Deep learning

**DOI:** 10.1101/2025.03.22.644762

**Authors:** Manolo F. Perez, Ricarda Riina, Brant C. Faircloth, Marcelo B. Cioffi, Isabel Sanmartín

## Abstract

Recognizing species boundaries in complex speciation scenarios, including those involving gene flow and demographic fluctuations, remains a challenge, particularly given the diversity of existing species concepts. Promising recent approaches adopt an integrative taxonomy that combines multiple sources of evidence (e.g., genetic, morphology, geographic distributions), reflecting different properties associated with the dynamics of the speciation continuum. The use of statistical inference methods for model comparison, such as approximate Bayesian computation, approximate likelihood approaches, and machine learning, has improved the better assessment of species boundaries in such contexts. However, most existing approaches involve analyzing genetic and phenotypic/geographical information separately, followed by visual/qualitative comparison. Methods that integrate genetic information with other sources of evidence remain limited to simple evolutionary models and are typically unable to analyze more than a few hundred loci across a maximum of a few tens of samples. Here, we present a deep learning approach (DeepID) that combines two convolutional neural networks to integrate genomic data (thousands of loci or single nucleotide polymorphisms, SNPs) and trait information into a unified framework. Using both simulated and empirical data sets, we evaluate the power and accuracy of this approach for discriminating among competing divergence speciation scenarios (with minimal ongoing gene flow) across a varying number of SNPs and traits, as well as different levels of missing data. Analyses based on genomic or trait data alone yielded a slight lower accuracy, whereas integrating genomic and trait data resulted in improved performance. When we violated the speciation model by including extensive migration, approaches incorporating trait data were less affected than those relying solely on genomic information. Together, these results suggest that combining genomic and trait data may capture complementary signals associated with different stages of the speciation process. Moreover, our approach successfully recovered the expected delimitation scenarios in empirical data sets from a plant (*Euphorbia balsamifera*) and a fish (*Lepomis megalotis*) species complex. We argue that our method is a flexible and promising approach, allowing for complex scenario comparison and the use of multiple types of data. Combining genomic and trait data likely captures complementary signals associated with different stages of the speciation process, reflecting the fact that speciation is a continuum in which genetic and phenotypic divergence may proceed at different rates.

## INTRODUCTION

Assessing species boundaries is a challenge that has been the focus of significant conceptual (de Queiroz 2007 and references therein) and methodological debate (see Carstens et al. 2013 for a review). The most promising species delimitation approaches suggest that a taxonomic framework integrating multiple sources of evidence, such as genetic, phenotypic (continuous and discrete traits), and ecotypic (geographic distribution, environmental preferences) data, can recover information from diverse stages of the speciation continuum (Carstens et al. 2013; Solís-Lemus et al. 2015; Eberle et al. 2018; Wibberg et al. 2021). Most strategies to combine these different lines of evidence consist of analyzing the component parts, followed by a comparison step that combines them by congruence or cumulation (sensu Padial et al. 2010); a standardized data format has recently been developed to facilitate these efforts (Miralles et al. 2022). Although this type of approach has made combining delimitation results from different data sources easier, the problem remains to analyze multiple sources of evidence in an integrative framework. iBPP (Solís-Lemus et al. 2015) is among the few methods that take an integrative approach by modifying the multi-species coalescent (MSC) delimitation method implemented in BPP (Yang 2015) to accommodate continuous traits. The integrated analysis of trait and genetic data showed promising results (Solís-Lemus et al. 2015), although BPP-based approaches like iBPP are currently limited to analyzing data from a few hundred loci collected across a modest number of samples (Oliveira et al. 2015; Eberle et al. 2019; Rabiee and Mirarab 2021). Moreover, because iBPP and BPP are based on the MSC model, they are open to the criticism of species over-splitting: they may recover population genetic structure rather than species limits, even in the presence of ongoing gene flow (Sukumaran and Knowles 2017; Chambers and Hillis 2020).

To reduce the effects of over-splitting, Jackson et al. (2017) suggested incorporating the genealogical divergence index (*gdi*) into strategies that estimate species limits, and subsequent work showed that its inclusion in BPP inferences improves accuracy while reducing over-splitting (Leaché et al. 2019). However, several drawbacks of this heuristic metric have been identified in recent studies. For example, *gdi* showed a high sensitivity to biased and/or incomplete geographic sampling when coupled with limited dispersal. In these cases, limited dispersal can create a genetic diversity gradient that causes population structure to be interpreted as species limits, especially when taxa are insufficiently sampled (Hausdorf and Hennig 2020; Chambers and Hillis 2020; Mason et al. 2020). Additionally, the large interval of *gdi* values for which delimitation is ambiguous (0.2–0.7), the fact that *gdi* does not take into account different population sizes among the groups being delimited (Rannala and Yang 2020), and the observation that *gdi* is sensitive to gene flow (Jiao and Yang 2021) have also been criticized. As an alternative, Rannala and Yang (2020) suggested considering intra-and interspecific limits for the (1) migration rate and (2) absolute divergence time in generations. They also introduced (3) an alternative heuristic metric – the gene tree probabilities *P*_a_ and *P*_b_; their equation 7 – which provides an approach similar to *gdi* while accounting for differences in population size among putative species (Rannala and Yang 2020).

Another strategy used to tackle the over-splitting issue in species delimitation is to model the population-level processes leading to speciation during inference (Sukumaran and Knowles 2017; Smith and Carstens 2020). These approaches can be classified into two main frameworks. The first consists of reference-based methods, which analyze groups of organisms with unknown species boundaries by using information either from closely related and well-delineated populations/species or from a subset of the same dataset being analyzed consisting of individuals with “undisputed” intraspecific and interspecific limits. The methodological strategies used to create these reference-based approaches involve *gdi* values (Leaché et al. 2021), estimating parameters associated with the protracted speciation process (Sukumaran et al. 2021), or supervised machine learning (Derkarabetian et al. 2022). However, the expert knowledge and natural history information needed to identify well-delineated or “undisputed” groups are often unavailable, which can limit the utility of these approaches in many taxa (Bebber et al. 2014; Wheeler 2014; Costello et al. 2020). The second suite of approaches that incorporate population-level processes is based on explicit modeling of demographic inference. These methods rely on genetic data sets simulated under competing speciation scenarios using a coalescent framework, and then compare the simulated data with the empirical data to select the speciation scenario that best fits the empirical data. The advantage of these methods is that they can be used in groups for which we lack expert knowledge of taxonomic limits. They also contribute to our understanding of the speciation process, especially for taxa with complex demographic histories, by explicitly incorporating events such as population size fluctuations or gene flow, thereby reducing incorrect delimitations arising from model violations (Jackson et al. 2017; Perez et al. 2022). Examples of this second suite of approaches include approximate Bayesian computation (ABC; Camargo et al. 2012), approximate likelihood analysis (Morales et al. 2017), coupling ABC with a random forest classifier (Smith and Carstens 2020), and machine learning methods (see Salles and Domingos 2025 for a review on machine learning methods for species delimitation), including support vector machines (Pei et al. 2018) and convolutional neural networks on alignment matrices (Perez et al. 2022).

Despite these advancements, a primary challenge remains: existing integrative methods often suffer from computational bottlenecks that limit their use to small data sets, whereas more scalable heuristic or simulation-based approaches typically frequently rely on a single data type. To address these limitations, we introduce an integrated species delimitation approach, named DeepID, which combines key features of some of the methods described above. DeepID overcomes the computational constraints of MSC-based integrative methods (e.g., iBPP) by utilizing a neural network architecture capable of processing thousands of loci alongside phenotypic data. Furthermore, by training the model on simulations explicitly parameterized using *P*_a_ and *P*_b_, as suggested by Rannala and Yang (2020), DeepID inherently accounts for the “gray zone” of speciation. This approach reduces the risk of over-splitting associated with traditional MSC models and the sensitivity to biased sampling associated with previous heuristic metrics like *gdi*. More technically, DeepID is a simulation-based, supervised deep learning approach that integrates multiple sources of evidence, including genomic and trait (continuous or discrete) data, to explicitly model intra- and interspecific divergence events and demographic processes under alternative species delimitation hypotheses. We assess the accuracy of DeepID for discriminating among alternative species delimitation hypotheses by simulating genetic and trait data sets under divergence speciation scenarios with minimal ongoing gene flow and different numbers of species. We also evaluate the effect of including varying levels of gene flow and missing data. Finally, we demonstrate the utility of DeepID when applied to real biological systems by analyzing phylogenomic and trait data sets from two empirical case studies: a plant (sweet tabaiba (*Euphorbia balsamifera*); Riina et al. 2021) and an animal (longear sunfish (*Lepomis megalotis*); Kim et al. 2022).

## MATERIALS AND METHODS

### Overview

Supervised deep learning approaches consist of several steps that include model design followed by model training, validation, and testing (with simulated data) and/or model prediction (with empirical data). Supervised deep learning approaches can also take many forms, and the one that we selected is built upon the idea of simulation-based deep learning (Borowiec et al. 2022; Korfmann et al. 2023). Unlike standard supervised learning, which relies on large, manually labeled empirical data sets (e.g., “undisputed” species boundaries that are often unavailable in non-model taxa), simulation-based deep learning uses a simulator to create an exhaustive range of “ground-truth” training samples. We selected this approach because it allows us to explicitly define the biological boundaries of our classes (via *P*_a_ and *P*_b_ thresholds) and ensures the model is trained on the demographic complexities (e.g., high gene flow levels and differences in population sizes) relevant to the empirical systems under study. In practice, this means a deep-learning model is designed to classify alternative speciation scenarios; simulated or empirical data are used to parameterize these scenarios and generate training data sets that are then used to improve the classification ability of the deep learning model. The trained model is subsequently either (a) evaluated for accuracy using independent simulated data or (b) applied to empirical data to infer the speciation scenario that most likely generated the empirical data.

At a high level, our supervised deep learning model (DeepID) integrates two neural networks to identify the most likely speciation scenario – one convolutional neural network (CNN) for analyzing sequence data and a second CNN for analyzing trait data. A CNN is an artificial neural network with layers that perform convolutions to extract spatial features from the input, which can be used to discriminate meaningful dependency patterns among segregating/polymorphic sites (Perez et al. 2022) or, as applied here, to also extract information from continuous/discrete traits. In DeepID, the two independent CNNs are then integrated using a (late) fusion approach that automatically optimizes the weight or “importance” of each data type (SNPs and traits) to maximize the accuracy of the overall model. The output of this fusion layer is then fed into a multi-layer perceptron (MLP; Haykin 1999), a simpler architecture with fully connected layers that extracts hierarchical features from the input and connects them to the output classification layer (the underlying speciation scenario).

After designing the model for DeepID, we followed two general approaches for training and testing whether DeepID accurately classifies speciation scenarios. In the first, we trained and validated DeepID using different sets of fully simulated data, and we used additional simulations from each of the scenarios to determine the accuracy of our approach. In the second, we used empirical estimates from two different case studies (Riina et al. 2021; Kim et al. 2022; described below) to parameterize simulations. These simulated data sets were then used to train and validate case-specific versions of DeepID, which were subsequently applied to classify the speciation scenario most likely to have generated each empirical dataset.

### Model Design

We designed DeepID in Keras v2.10.0 with a TensorFlow v2.9.0 backend using Python v3.8.10. DeepID builds on the approach proposed in Perez et al. (2022) but differs in a key aspect: it integrates genetic data (as in Perez et al. 2022) with trait data (this study) within a unified deep learning framework to predict the most likely speciation scenario (among a set of competing scenarios) giving rise to the input data types.

We encoded genetic data (SNPs) as arrays using NumPy v1.21.2 where loci were represented by rows, individuals were represented by columns, and genotypes were coded as -1 for the major state and 1 for the minor state. We loaded the resulting arrays into a CNN, using the architecture described in Perez et al. (2022) with some modifications: (i) we implemented three 1D-convolutional layers, each with a kernel size of three and 256 neurons (Fig. 1); (ii) these layers were intercalated with batch normalization, following Sanchez et al. (2021); and (iii) we replaced average pooling with max pooling (Nikanjam and Khomh 2021). After the pooling step, the features were flattened (i.e., collapsed into a single linear vector) and fed to a MLP containing two fully connected (FC) layers with 128 neurons each and an additional FC layer containing 64 neurons (Fig. 1), intercalated with 50% dropout to avoid overfitting (Hinton et al. 2012).

**Figure 1.**
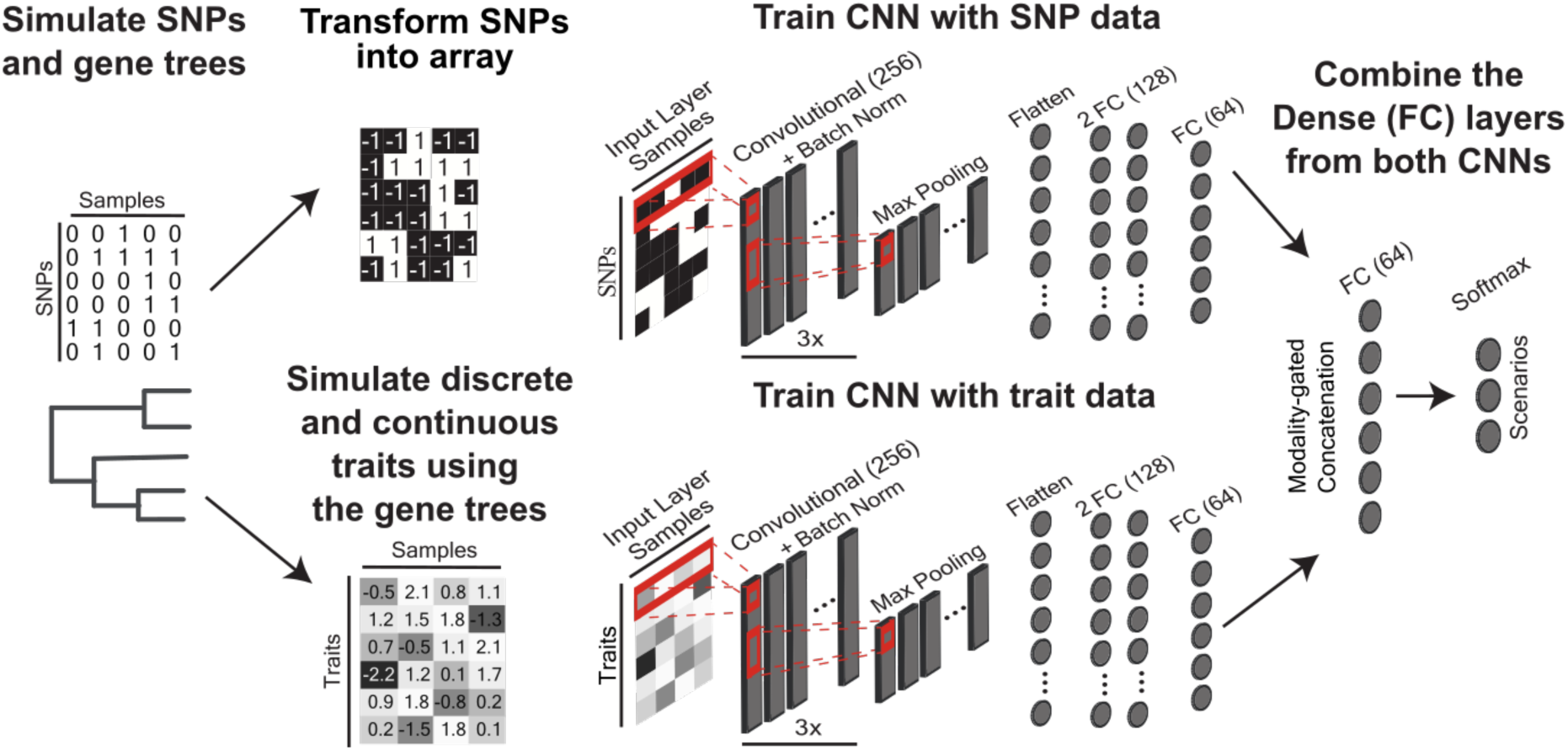
Outline of our deep learning species delimitation approach (DeepID) that combines two convolutional neural networks (CNN), one for the genetic data and one for the trait data. The first step of the process is to simulate genetic data (SNPs) from competing speciation scenarios and generate a SNP data matrix and associated phylogenetic trees for each simulation. Then, the SNP data are fed into one CNN, while the trees are used to simulate discrete and continuous traits that are input into a second CNN. Finally, DeepID combines the outputs of both networks using modality-gated concatenation and uses a softmax layer to calculate the probability of each delimitation scenario.

We also encoded trait data as NumPy arrays where traits were represented by rows and individuals were represented by columns. These data were then input into a CNN identical to that used for the genetic data. To enable integration of both networks, we added a final FC layer with 64 neurons (Fig. 1), matching the size of the last layer in the genetic-data network.

Finally, DeepID combines the final FC layers of the networks for each data type using a Modality-Gated Concatenation layer (inspired by Arevalo et al. 2017). Unlike standard concatenation, which assumes each data modality contributes equally to the final classification, this strategy introduces a trainable gating parameter for each data branch (i,e. one for the genetic data and one for the trait data). These gating parameters act as attention weights that scale the feature activations of each branch before they are merged. To avoid a priori bias toward a given data type, the gates are initialized with a 50/50 contribution for both branches (genetic data and traits). We designed this layer to be switchable, meaning the gating parameters can either be learned jointly with the network weights to maximize label accuracy or set to fixed, user-defined constants to test specific hypotheses regarding the importance of each data branch – meaning that it is possible to fix, for example, the contribution of the genetic data to 30% and the trait data to 70% if we want to give more weight to trait data in the final classification. If we set this layer to be learnable, the absolute values of these learned weights after training provide a direct metric of branch importance, allowing us to quantify the relative contribution of genetic data versus trait data in the species delimitation. We connected the concatenation layer to an output classification layer to calculate model probabilities using a softmax function (Fig. 1). We optimized the design of DeepID (number of FC layers, number of neurons per FC layer, etc.) by performing several test runs of the model using small subsets of simulated data which are described in the next section.

### Simulated Data: Generation

To evaluate the performance of DeepID, we designed a procedure to simulate data where we created three simple speciation scenarios involving divergence under minimal gene flow on a dataset containing six populations belonging to one, two or three species (Fig. 2). Then, we used a custom Python script that integrated *ms* v2018.11.17 (Hudson 2002) to simulate data sets under each of the three scenarios, and we labeled each set of simulated data with the name of the generative speciation scenario. Each simulation explicitly modeled intra- and interspecific divergence events (*τ*) using *P*_a_ and *P*_b_ (Rannala and Yang 2020) as a heuristic alternative to *gdi*. We calculated the relationship between τ and each lineage-specific *P* values (*P*_a_ and *P*_b_) by computing 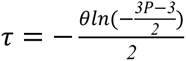 (derived from equation 7 in Rannala and Yang 2020), and we converted these divergence times to 4*τ*/*θ* (4*N*_e_ generations) time units as required by *ms*.

**Figure 2.**
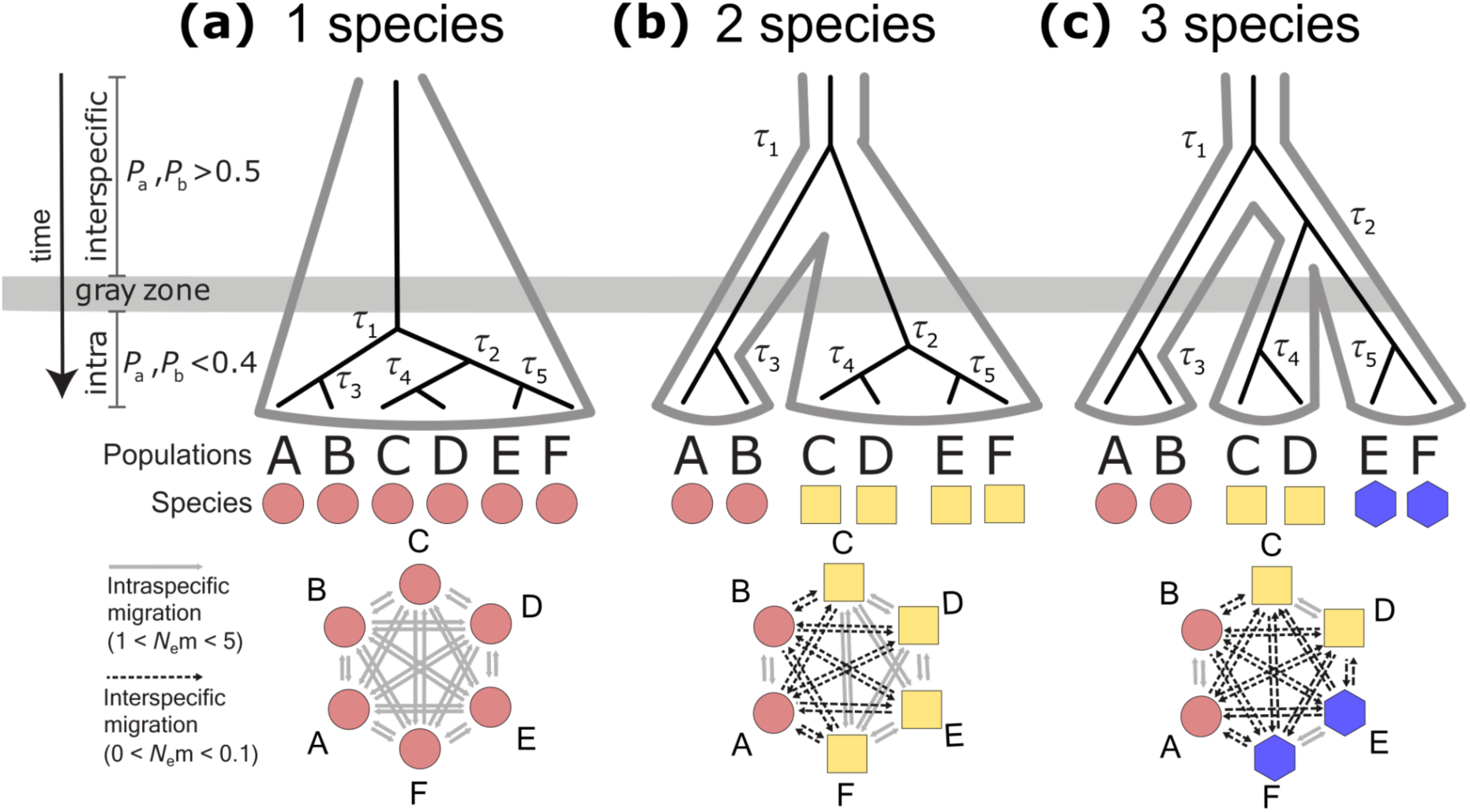
Schematic representation of the three simple speciation scenarios simulated, where the species tree is outlined in dark gray and the divergence events (*τ*) are depicted by the guide tree in black for six populations (A-F). Threshold levels for interspecific (*P*_a_ and *P*_b_ > 0.5) and intraspecific (*P*_a_ and *P*_b_ < 0.4) divergence events, as well as the uncertain interval (gray zone; 0.4 ≥ {*P*_a,_*P*_b_} ≥ 0.5) for species delimitation, were established according to Rannala and Yang (2020). In the scenario with only one species (a), *τ*_1_ (associated with the deepest node) was sampled from a uniform distribution with values that ensure *P*_a_ and *P*_b_ < 0.4, corresponding to an intraspecific divergence event, and all other *τ* values were simulated as more recent than *τ*_1_. In the two species scenario (b), the interspecific value of *τ*_1_ (associated with the deepest node) was sampled from a uniform distribution with values that ensure that *P*_a_ and *P*_b_ fell between 0.5 and 1, representing an interspecific divergence event, while all other *τ* values were sampled from a uniform distribution that renders *P*_a_ and *P*_b_ below 0.4, representing intraspecific divergence events. For the three species scenario (c), the interspecific nodes *τ*_1_ and *τ*_2_ were simulated with *P*_a_ and *P*_b_ values between 0.5 and 1 while ensuring that *τ*_1_ > *τ*_2_; all other intraspecific *τ* values (*τ*_3_, *τ*_4_, and *τ*_5_) were simulated as more recent than *τ*_2_. All scenarios included a baseline intraspecific (1 < *N*_e_m < 5) and interspecific (0 < *N*_e_m < 0.1) bidirectional asymmetric migration rate among relevant population pairs, which are depicted in the migration schemes associated with each speciation scenario. Throughout the figure, different colored symbols represent distinct species.

The simulation process began by setting the baseline mutation-scaled population size parameter (*θ* = 4*N*_e_μ) of each simulated population (A to F) to a value drawn from a uniform distribution that ranged from 0.0025 to 0.01. Then, we sampled five diploid individuals (10 haplotypes) from each population. For the speciation scenarios with more than one species (Fig. 2b, c), we also sampled a value for the first divergence time event (*τ*) from a uniform distribution spanning the minimum and maximum *τ*_1_ values required for both *P*_a_ and *P*_b_ (the two daughter lineages) to be within the interval of interspecific events (0.5 and 1), given their respective effective population sizes (measured by their corresponding *θ* value). If a speciation scenario included a second interspecific divergence event (Fig. 2c), we sampled a second divergence time *τ*_2_ from a uniform distribution where the minimum and maximum values would also ensure that *P*_a_ and *P*_b_ were higher than the interspecific threshold (0.5) and also that *τ*_2_ < *τ*_1_. Similarly, we sampled values of *τ*_3_ from a uniform distribution spanning minimum and maximum values that would ensure both *P*_a_ and *P*_b_ were within the intraspecific thresholds (between 1/3 and 0.4). We simulated the remaining times for population divergence events (*τ*_4_ and *τ*_5_) in 4*N*_e_ generational units by drawing values from a uniform distribution, where more recent values were drawn from a range that was constrained to be younger than the subtending intraspecific divergence event (Fig. 2). We did not sample *τ* values that would result in *P*_a_ and *P*_b_ values between 0.4 and 0.5 because these would be regarded as ambiguous and fall into a gray zone (Rannala and Yang 2020) for species delimitation (Fig. 2). Apart from recommendations for *P*_a_ and *P*_b_ thresholds, Rannala and Yang (2020) suggest limits on gene flow and divergence time (in number of generations) to distinguish species from populations. Following these guidelines, all DeepID speciation scenarios included a baseline intraspecific (*N*_e_m between 1 and 5) and interspecific (*N*_e_m < 0.1) migration rate among relevant population pairs. While Rannala and Yang (2020) also propose divergence time thresholds (< 10^3^ generations for intraspecific and > 10^4^ generations for interspecific), we decided to not include these temporal thresholds in our simulation to make the DeepID approach more flexible.

After setting these parameters, we input them to *ms* to simulate 1,000 independent loci for samples in each population under the Wright-Fisher neutral model where we output a fixed number (n = 1) of segregating sites and the gene tree (-T option) for each locus. For each iteration, we saved the model parameters, retained segregating sites as an array containing 1,000 loci, and randomly sampled and retained 100 topologies (the number of traits to be simulated, described below) from the larger set of 1,000 gene trees output by *ms*. We repeated this process 11,000 times for each speciation scenario.

After running the *ms* simulations, we input each set of 100 retained topologies from each simulation (n = 11,000) across all speciation scenarios to phytools v0.7 (Revell 2012) to generate simulation-specific trait data sets. Since traits can be both discrete and continuous, and because continuous traits can be simulated under different models, we repeated this process three times to simulate: 100 discrete traits with three states using the sim.history command (Markov k-model - Mk) with symmetric rates, 100 continuous traits under the Brownian-Motion (BM; sig2 = 0.06) model, and 100 continuous traits under the Ornstein-Uhlenbeck (OU; sig2 = 0.06, alpha = 0.2, theta = 0) model. The end result of this process created three distinct groupings of genetic+trait data: (1) genetic data simulated under each speciation scenario plus a set of discrete traits, (2) genetic data simulated under each speciation scenario plus a set of continuous traits (BM Model), and (3) genetic data simulated under each speciation scenario plus a set of continuous traits (OU Model).

### Simulated Data: Model Training, Validation, and Testing

Although DeepID is capable of analyzing mixtures of trait data, we trained, validated, and tested the model for each distinct group of data separately (genetic data+discrete traits, genetic data+continuous (BM) traits, genetic data+continuous (OU) traits). Within each of these major categories, we combined the labeled simulations derived from each speciation scenario, randomly shuffled the data using the train_test_split function in Scikit-learn v1.2.0, and divided the data into a portion used to fit the model (the “training set”, n = 7,500 simulations for each scenario), a portion used to evaluate the training procedure and tune the artificial neural network parameters (the “validation set”; n = 2,500 simulations for each scenario), and a portion used to evaluate the accuracy of the model (the “test set”; n = 1,000 simulations for each scenario). Before analyzing data sets that included continuous traits (genetic data+continuous (BM) traits, genetic data+continuous (OU) traits) and because trait data are often recorded using different units of measure, we standardized the trait values by scaling trait values to unit variance using the StandardScaler function in Scikit-learn.

To speed the training process and avoid memory overflows during training, smaller batches (minibatches) of simulated data can be input to the deep learning model in parallel. An “epoch” refers to each time the entire dataset is passed through the model. We input the simulation data to DeepID in mini-batches containing 250 simulations for each of the 100 epochs. We evaluated the training procedure on the validation set using a categorical cross-entropy loss function and employed rectified linear units (ReLU; Nair and Hinton 2010) as the non-linear activation functions, as they help mitigate the vanishing gradient problem and facilitate efficient backpropagation of the loss signal. This approach is commonly used in the context of population genetics and phylogeography for classification problems based on SNP data (Flagel et al., 2019; Fonseca et al., 2021; Perez et al., 2022). We also optimized the network parameters with stochastic gradient descent using a learning rate of 0.001, a momentum coefficient of 0.9, and the Nesterov accelerated gradient method (Sutskever et al. 2013). To avoid overfitting, we saved only the epoch with the highest accuracy in the validation set using model checkpointing.

After we trained DeepID with labeled data sets, we used a third set of simulated, labeled data, that was not used for training or validation (the “test set”; n = 1,000 simulations) to evaluate the accuracy of the network. Specifically, we input the test set to DeepID and compared the predicted speciation scenario with the true labels of the simulated data to build a confusion matrix of predicted versus generative scenarios using a cross-validation step. Accuracy was measured as the percentage of correct assignments for the scenarios from which the data were simulated.

### Simulated Data: Varying Number of SNPs and Traits

Different empirical studies sample different numbers of SNPs and traits from populations. To assess how these differences affected the ability of DeepID to identify the generative speciation scenario, we used NumPy to create random subsamples of the simulated data to evaluate the effects of varying the numbers of SNPs (10, 50, 1000) and traits (10, 50, 100) on model accuracy. Because some studies collect only SNP or only trait data, we also generated two additional data sets consisting exclusively of SNP data or trait data. After creating these additional data sets, we trained, validated, and tested DeepID using the same general procedures described above.

### Simulated Data: Assessing the Impact of Missing Data

Empirical data often have missing entries for genetic and/or trait data (Ahrens et al. 2016), and we were interested in testing how varying levels of missing data would affect our approach. A typical combined dataset may include two forms of missing data: absent values for loci or traits scattered across sampled individuals, and unequal sampling of individuals among data types. To emulate this, we modified copies of the simulated data in two ways: (i) by removing values from random cells across SNP and trait simulated data sets (missing values), and (ii) by removing all cells from a random subset of simulated individuals (unequal sampling). Specifically, we used NumPy to randomly insert missing data into the SNP matrices at varying levels (20%, 40%, and 60%), either by removing individual cells or entire individuals, and encoded missing values as 0 (Kirschner et al. 2022). Similarly, we modified the trait data matrices by inserting 0 values at the same percentages. After creating these additional data sets, we trained, validated, and tested DeepID using the procedures described above.

### Simulated Data: Speciation Scenarios with Migration

Methods based on the MSC model can be negatively affected by gene flow (Jackson et al. 2017; Sukumaran and Knowles 2017; Leaché et al. 2019), so we explored how post-divergence gene flow among populations influences the performance of DeepID. Hence, we modified the baseline simulation code for speciation scenarios with two (scenario b; Fig. 2) and three species (scenario c; Fig. 2) to include continuous bidirectional migration among all species pairs (AB `↔` CDEF in scenario 2b; AB `↔` CD `↔` EF in scenario 2c) at higher levels than the original scenarios (*N*_e_m < 0.1), beginning at their interspecific divergence and continuing to the present (using the -ma option in *ms*). Because rates of post-divergence gene flow can vary (Jackson et al. 2017), we assessed how different migration rates affected the performance of DeepID. So, we simulated a combined genetic (1,000 SNPs) and trait data (100 traits using discrete, BM, or OU models) using three different migration rates (*N*_e_m = 0.125, 0.25, and 0.5). This produced three distinct groupings of genetic+trait data, as before (genetic data+discrete traits, genetic data+continuous (BM) traits, genetic data+continuous (OU) traits) at three different migration rates. We trained, validated, and tested nine versions of DeepID – corresponding to each combination of genetic+trait data and migration rate – using the procedures described above.

We were also interested to know how well DeepID performed when it was trained with one class of data but used to classify another (model misspecification), so we ran two additional tests: (1) classifying populations simulated with high migration using DeepID models trained on populations only with baseline migration, and (2) classifying populations simulated with baseline migration using models trained on populations exhibiting high migration.

### Empirical Data: Overview

Although simulations are often used to train, validate, and test deep learning models, the classification problems in which we are ultimately interested involve empirical data. We selected two case studies involving one plant (sweet tabaiba (*Euphorbia balsamifera*); Riina et al. 2021) and one animal (longear sunfish (*Lepomis megalotis*); Kim et al. 2022) species complex in order to demonstrate how DeepID performs with more complex data from real empirical systems. These studies are examples of long-standing taxonomic controversies that have recently been addressed with phylogenomic data and trait-based evidence, resulting in new proposals regarding species limits (Riina et al. 2021; Kim et al. 2022). Each study also represents a speciation scenario with different levels of complexity, from simple divergence with no secondary contact (sweet tabaiba; Riina et al. 2021) to speciation followed by frequent hybridization and introgression (longear sunfish; Kim et al. 2022). Both studies also contain different proportions and patterns of missing data, which may represent empirical systems more accurately than our simulated scenarios. Finally, each study used a different method to delimit species: comparative integrative taxonomy (sweet tabaiba; Riina et al. 2021) versus statistical species delimitation (longear sunfish; Kim et al. 2022). Comparing the results from DeepID with the delimitation hypotheses proposed in the original studies (Riina et al. 2021; Kim et al. 2022) allowed us to explore the power and limits of the approach we developed.

### Empirical Data: Sweet Tabaiba

The *Euphorbia balsamifera* or “sweet tabaiba” complex (*Euphorbia balsamifera* s.l., Riina et al. 2021) belongs to the *Euphorbia* (Euphorbiaceae) subgenus *Athymalus* (Peirson et al. 2013) and has long been a taxonomic riddle (Riina et al. 2021). The species complex currently includes three taxa, *E. adenensis* Deflers, *E. balsamifera* Ait., and *E. sepium* N.E.Br. (Riina et al. 2021), although the taxonomic rank and count of members within the group have shifted over time with the use of different delimitation criteria.

Historically, several species have been associated with what is currently referred to as the species complex (Riina et al. 2021) including *Euphorbia balsamifera* Ait. (Aiton 1789), *E. adenensis* Deflers (Deflers 1887), *E. sepium* N.E.Br. (Brown et al. 1911), *E. rogeri* N.E.Br. (Brown et al. 1911), and *E. capazii* Caball. (Caballero 1935). Changes to this taxonomy began with Maire (1938) who combined *E. rogeri* and *E. capazii* to form *E. b.* var. *rogeri* and who also subsumed *E. sepium* into *E. b.* subsp. *sepium*. Bally (1965) subsequently noted that *E. adenensis* appeared quite similar to *E. balsamifera*, despite their range disjunction, so he refined the taxonomy by naming *E. adenensis* as a subspecies of *E. balsamifera*: *E. b.* subsp. *adenensis*. Govaerts et al. (2000) later subsumed *E. b.* subsp. *sepium* under *E. b.* subsp. *balsamifera*, while Molero et al. (2002) resurrected *E. b.* subsp. *sepium* based on karyological data. However, the taxonomy of Govaerts et al. (2000) prevailed and was adopted by Peirson et al. (2013) in their molecular study using one nuclear and one chloroplast locus. More recently, Villaverde et al. (2018) used DNA sequence data of 296 exons collected from a large sample of individuals across the distribution of the *E. balsamifera* complex to show that the group consisted of three well-supported lineages, with *E. b.* subsp. *sepium* sister to a clade formed by the geographically disjunct *E. b.* subsp. *balsamifera* and *E. b.* subsp. *adenensis*. Finally, Riina et al. (2021) employed an integrative taxonomic approach combining the results from Villaverde et al. (2018) with morphology data (leaf measurements), climatic niche analysis, and geographic occurrence data (i.e., disjunct distributions) to elevate *E. b.* subsp. *sepium, E. b.* subsp. *balsamifera*, and *E. b.* subsp. *adenensis* to the rank of species (*E. sepium, E. balsamifera*, *E. adenensis*). However, a recent study (Thulin et al. 2025) has challenged the reinstatement of *E. balsamifera* and *E. adenensis* as distinct species and instead proposed reverting to the previous taxonomy of two geographically disjunct subspecies within *E. balsamifera*.

Given the somewhat muddled history of this plant group, we used DeepID to explicitly test which of five different speciation scenarios (Fig. 3a) is the most likely given the data. Specifically, we tested whether the sweet tabaiba complex consists of: (1) a single species containing all three taxa as three subspecies or varieties (Fig. 3a.i; Molero et al. 2002; Villaverde et al. 2018), (2) two species, with *E. adenensis* sister to a branch including both *E. b.* subsp. *sepium* and *E. b.* subsp. *balsamifera* (Fig. 3a.ii); (3) two species, with *E. sepium* sister to a branch containing both *E. b.* subsp. *adenensis* and *E. b.* subsp. *balsamifera* (Fig. 3a.iii; Thulin et al. 2025); (4) three distinct species, with *E. sepium* sister to a clade containing *E. balsamifera* and *E. adenensis* (Fig. 3a.iv; Riina et al. 2021); and (5) the same three species and topology as in (4), but with gene flow higher than the baseline (*N*_e_m < 0.1) occurring among all species. Because there is no evidence of contemporary gene flow among the three species, we simulated a more realistic ancient migration scenario, in which these higher levels of interspecific migration were allowed only during an initial time interval, from the time of species divergence (*τ*_1_ and *τ*_2_) up to half of the total divergence time (*τ*_1_/2 and *τ*_2_/2, respectively; Fig. 3a.v). Migration rates (*N*_e_m) for this interval of higher migration were drawn from a uniform distribution *U*(0.1, 0.5).

**Figure 3.**
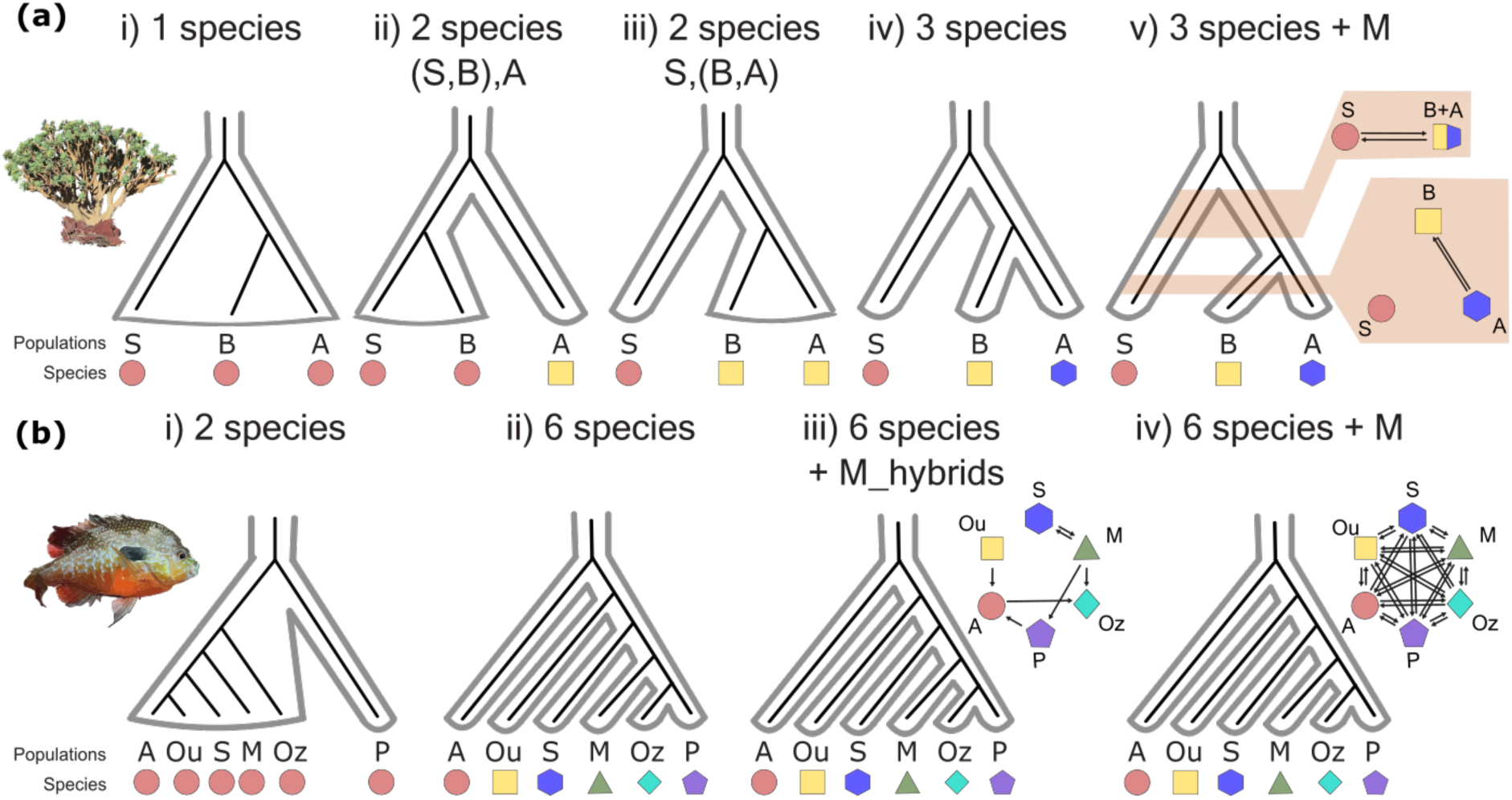
Schematic representation of the simulated species delimitation scenarios in (a) the sweet tabaiba (*Euphorbia balsamifera* species complex) and (b) the longear sunfish (*Lepomis megalotis* species complex). The species tree topology for each scenario is outlined in gray, with the relationship among populations depicted by the guide tree in black. We used the same baseline levels of *P*_a_ and *P*_b_ shown in Fig. 2 to define intra- and interspecific divergence times, and we used the baseline intraspecific (1 < *N*_e_m < 5) and interspecific (0 < *N*_e_m < 0.1) migration levels from Fig. 2 in all scenarios. Increased levels of gene flow (higher interspecific migration rates, 0.1< *N*_e_m < 0.5) are depicted as black arrows in the migration scheme shown on the right side of each relevant scenario (b.iii, b.iv, and a.v - with periods of increased gene flow, i.e. from the initial split of two lineages from a common ancestor to halfway through the total divergence time, and their respective migration schemes are represented with a light orange outline in the latter). We used different symbols to represent distinct species and labeled those symbols with the species or population name. In panel (a): S = *E. sepium*, B = *E. balsamifera*, and A = *E. adenensis* (corresponding to *E. b.* subsp. *sepium*, *E. b.* subsp. *balsamifera* and *E. b.* subsp. *adenensis*, respectively, in scenarios with one or two species). In panel (b): A = *L. aquilensis*, Ou = *L*. sp. Ouachita, S = *L. solis*, M = *L. megalotis*, Oz = *L*. sp. Ozark, P = *L. peltastes*. Species were illustrated by one of the authors (MFP) in Adobe Illustrator CC 2018.

For empirical data, we used the original 428 locus DNA sequence matrix from Villaverde et al. (2018), along with data from four continuous traits related to leaf size and shape (leaf maximum length, leaf maximum width, length from the leaf base to the point of leaf maximum width, and mucron length) used by Riina et al. (2021). Each individual in the sequence matrix was represented by a consensus sequence, and we used a custom script that integrated fasta2genlight from adegenet v2.1.7 (Jombart 2008) to identify variable positions in each alignment and retain the single SNP from each locus having the fewest missing values. Missing data by site averaged 17.7%.

Based on these empirical data, we generated sets of simulated data for model training and validation following procedures and parameters (i.e., 11,000 simulations) similar to those described above (**Simulated Data: Generation**), except that: (a) we used a different set of speciation scenarios (Fig. 3a), (b) we simulated a number of SNPs equal to the number in the empirical data (428), and (c) some simulation parameters were based on empirical estimates derived from *Euphorbia* populations. Specifically, we sampled *θ* values from a uniform distribution that ranged from 0.0005 to 0.006, the limits of which we derived by dividing the upper and lower estimates of *θ* (ca. 2-5) from different populations by the average length of the sequence of ∼1700 bp (Table 1 in Rincón-Barrado et al. 2024), and then allowed species-specific *θ* values to vary between 0.5 and 2 times these baseline values. We used these *θ* values to calculate intra- and inter-specific divergence times for each speciation scenario (Fig. 3a) using *P*_a_ and *P*_b_, and we input the sampled values to *ms* to simulate SNP data sets that were the same size as the empirical data (i.e., the same number of individuals included for each species and 428 SNPs; Table 1). We randomly sampled and retained 4 topologies of the 428 gene trees output by *ms,* which we used to simulate continuous trait data using the BM and the OU models in phytools. For the simulated SNPs, we converted the major state to -1, the minor state to 1. We incorporated missing values into the simulated data using a procedure similar to the one described above (**Simulated data: Assessing the Impact of Missing Data**) by randomly inserting missing values (0) at a proportion (17.7%) similar to that of the empirical data. We trained DeepID using the simulated data as described in the **Simulated Data: Model Training, Validation, and Testing** section.

**Table 1.** Sample sizes for each putative species in the two empirical data sets. N_SNPs = number of samples containing SNP data. N_traits = number of samples containing trait data.

| Putative species | N_SNPs<sup>1</sup> | N_traits |
| --- | --- | --- |
| Sweet tabaiba |  |  |
| *E. sepium* | 10 | 29 |
| *E. adenensis* | 19 | 22 |
| *E. balsamifera* | 80 | 29 |
| Longear sunfish |  |  |
| *L. aquilensis* x *L. sp. Ouachita* | 4* | 0 |
| *L. aquilensis* x *L. peltastes* | 6 | 19 |
| *L. aquilensis* | 68 | 298 |
| *L. megalotis* x *L. solis* | 3* | 0 |
| *L. megalotis* | 44 | 204 |
| *L. sp. Ouachita* | 10 | 28 |
| *L. sp. Ozark* x *L. aquilensis* | 1* | 0 |
| *L. sp. Ozark* x *L. megalotis* | 8 | 79 |
| *L. sp. Ozark* | 15 | 76 |
| *L. peltastes* x *L. megalotis* | 10 | 34 |
| *L. peltastes* | 19 | 481 |
| *L. solis* x *L. megalotis* | 5 | 27 |
| *L. solis* | 36 | 229 |

\*Missing values were added for all traits in these samples. <sup>1</sup>The number of individuals used in our deep learning analyses was the same as N\_SNPs and the trait data were randomly subsampled to match N\_SNPs when N\_traits > N\_SNPs or completed with missing values when N\_traits < N\_SNPs.

DeepID assumes that simulated and empirical data sets have the same dimensions. However, as it is common in empirical data sets, the number of individuals in the sweet tabaiba data set for which SNP data were collected differed from the number of individuals for which trait data were available (Table 1). To obtain empirical data sets matching the dimensions of the simulations used to train DeepID, we applied two different approaches to generate 100 randomly resampled data sets combining the available trait data with the available SNP data from the same species. Specifically, for *E. sepium* and *E. adenensis*, which have more individuals sampled for traits (29 and 22, respectively) than for SNPs (10 and 19, respectively), we used NumPy to randomly subsample the trait dataset to match the number of individuals in the SNP matrix (Table 1). Conversely, for *E. balsamifera*, which has more individuals sampled for SNPs (80) than for traits (29), we used NumPy to randomly assign the suite of traits of one individual to another, random individual in the SNP matrix. For the remaining individuals that were not randomly assigned trait profiles, we simply coded their missing entries as 0 for all trait values. We repeated this process 100 times to maximize the use of the available data, minimize the potential for bias associated with random trait assignment, and obtain a distribution of posterior model probabilities rather than a single point estimate. We used the resulting 100 data sets (resampled from the available empirical data, as described above) as input to perform predictions with DeepID.

### Empirical Data: Longear Sunfish

The speciation scenarios presented for the *Euphorbia balsamifera* species complex (Fig. 3a) are relatively simple because the original studies postulate lineage divergence with no secondary contact. To assess the accuracy of DeepID with a more complex empirical example, we used data from the *Lepomis megalotis* (Rafinesque 1820) species complex. *Lepomis* (Centrarchidae, Percoidea) is a genus of freshwater fishes characterized by frequent hybridization, which has led to an unstable taxonomy (Bolnick 2009). Similar to the *E. balsamifera* complex, the *L. megalotis* complex has an intricate taxonomic history (see Kim et al. (2022) for a detailed description). It once included as many as 16 species (Bailey 1938; Gilbert 1998), which were later synonymized under *L. megalotis*. Several of these taxa were subsequently recognized as subspecies of *L. megalotis* (*L. m. megalotis*, *L. m. peltastes*, *L. m. aquilensis*, *L. m. breviceps*, and *L. m. occidentalis*). However, *L. m. peltastes* was later reinstated to species rank (*L. peltastes*; Bailey et al. 2004), despite limited supporting evidence (Kim et al. 2022). Kim et al. (2022) proposed that the complex comprises six relatively ancient lineages (*L. megalotis*, *L. peltastes*, *L. aquilensis*, *L. solis*, *L.* sp. Ouachita, and *L.* sp. Ozark), based on the combined results of *gdi* analysis and demographic (genetic) modeling. They further validated their findings using morphological data analyzed with a cross-validation linear discriminant analysis (LDA).

Given this complicated history, we were interested in using DeepID to discriminate between the following alternative species delimitation scenarios: 1) the *L. megalotis* complex consists of only two species (Fig. 3bi), following the traditional taxonomic arrangement (Bailey et al. 2004); 2) the *L. megalotis* complex consists of six species (Fig. 3bii), according to Kim et al. (2022); 3) the *L. megalotis* complex consists of six species with unidirectional gene flow between several pairs of species (Fig. 3biii), which were identified as potential hybrids by Kim et al. (2022), based on their integrative species demilitation results; and 4) the *L. megalotis* complex consists of six species with bidirectional gene flow between all species pairs (Fig. 3biv).

The empirical data sets we used consisted of 118,946 biallelic SNPs genotyped across 229 individuals (Table 1) as well as 28 continuous and 3 discrete traits collected from 1,475 individuals (Table 1; available from http://dx.doi.org/10.5061/dryad.dbrv15f05). Similar to the approach we used for *E. balsamifera*, we generated sets of simulated data for model training and validation following the procedures and parameters (i.e., 1,000 SNPs and 11,000 simulations) described above (**Simulated Data: Generation**), except that: (a) the set of speciation scenarios differed (Fig. 3b), and (b) some simulation parameters were based on empirical estimates from *Lepomis*. Specifically, we sampled *θ* from a uniform distribution ranging from 0.001 to 0.02, the limits of which were based on the range of *θ* values in Supplementary Figure 4 of Kim et al. (2022). We allowed species-specific *θ* to vary between 0.5 and 2 times these baseline values and we used the obtained *θ* values to calculate divergence times according to intra- and interspecific *P*_a_ and *P*_b_ thresholds for each speciation scenario. We input the sampled values to *ms* to simulate data sets of 1,000 SNPs that represented each speciation scenario. For the scenario presented in Figure 3b.iii, we also included unidirectional migration from *L*. sp. Ouachita to *L. aquilensis*, *L. peltastes* to *L. aquilensis*, *L. solis* to *L. megalotis*, *L. aquilensis* to *L*. sp. Ozark, *L. megalotis* to *L*. sp. Ozark, *L. megalotis* to *L. peltastes*, and *L. megalotis* to *L. solis* to match the predictions regarding potential hybrids of Kim et al. (2022). For scenario in 3biv, we modeled bidirectional migration among all six species pairs. In both scenarios with higher migration (Fig. 3b.iii and 3b.iv), we sampled migration rates (*N*_e_m) from a uniform distribution *U*(0.1, 0.5), which are higher than the *N*_e_m < 0.1 baseline. Similar to the procedure described above, we randomly sampled 31 topologies (matching the number of sampled traits) from the 1,000 gene trees generated by ms and used them as input to phytools to simulate three discrete traits and 28 continuous traits under the BM and OU models. Prior to training, we randomly added missing data to the simulated SNP data sets to mirror the proportion of missing SNPs (44.4%) observed in the empirical dataset, following the approach described in **Simulated data: Assessing the Impact of Missing Data**. We then trained the network as described above (**Simulated Data: Model Training, Validation, and Testing**).

Similar to the situation with the *Euphorbia balsamifera* complex, DeepID assumes that simulated and empirical data sets have the same dimensions. However, the empirical data for *Lepomis* are unbalanced, with SNP data available for 229 individuals and trait data for 1,475 individuals. To address this, we followed a similar resampling procedure to that described for the *Euphorbia* complex above, generating data sets containing both SNP and trait data for 229 individuals. Specifically, we used NumPy to randomly assign suites of traits to individuals, matching subspecies and hybrids where possible. For three potential hybrids (*L. aquilensis* x *L*. sp. Ouachita; *L. megalotis* x *L. solis*; *L*. sp. Ozark x *L. aquilensis*, Kim et al. 2022), no trait data were available, so we simply assigned them missing values for all traits in each resampled dataset. We repeated this process 100 times to thoroughly sample the available data, to minimize bias associated with random trait assignment, and to produce a posterior distribution of model probabilities rather than a single point estimate. We used the resulting 100 data sets (resampled from the available empirical data, as described above) as input to perform predictions with our trained network.

## RESULTS

### Model Design and Training

The most costly component of the DeepID approach is the training process because the inference time after the network is trained is almost instantaneous. For the larger simulated data sets (e.g., those containing 1,000 SNPs), training the network on an Nvidia L4 GPU took ∼30 minutes for both the combined and SNP-only models, while each trait-only model took ∼5 minutes to train. Running times were measured with the command time.time().

### Simulated Data: Varying Number of SNPs and Traits

DeepID exhibited high classification accuracy when differentiating among the test data sets simulated under the three simple speciation scenarios (Fig. 4). When analyzing the same number of traits and SNPs, we usually observed slightly lower levels of accuracy when DeepID attempted to classify speciation scenarios using only simulated discrete traits, while accuracy was similar when using only simulated continuous traits or only SNP data (Fig. 4a). When using a combination of simulated SNP+simulated trait data, the percentage of correct assignments to the simulated scenario was always higher than 95% for the combined data sets comprising 10 SNPs and always higher than 99% when we used 50 and 1,000 SNPs (Fig. 4b). We observed a general trend of increasing classification accuracy when we analyzed more characters, whether looking at increasing the number of continuous or discrete traits or increasing the number of SNP loci (Fig. 4). The proportional contribution of SNPs to the classification decisions – the learned weights given to the SNPs data branch in the gated concatenation – increased as more SNPs were used. Moreover, for a given number of SNPs, this proportional contribution of SNPs decreased as more traits were used in the combined data set (Fig. 4b).

**Figure 4.**
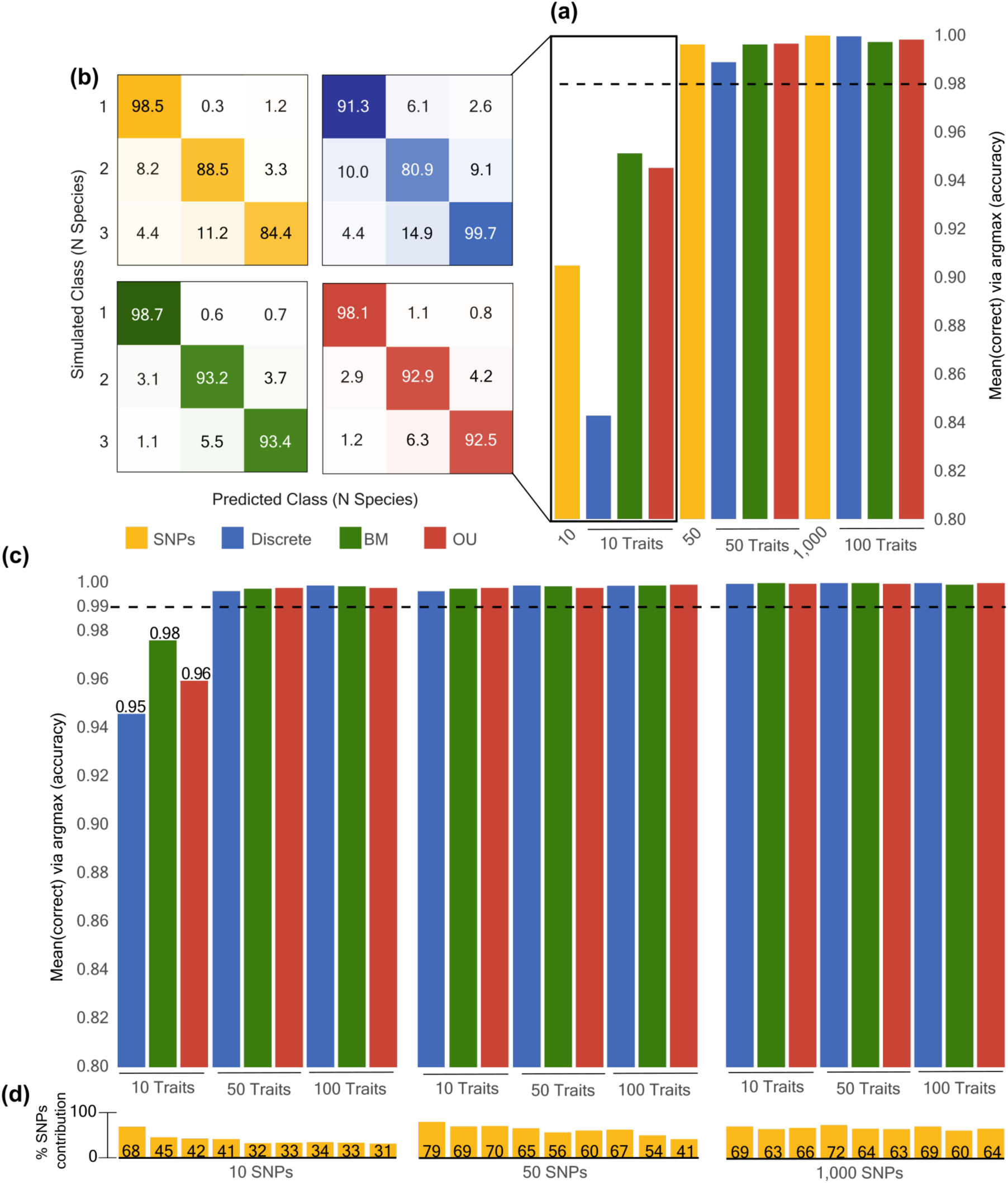
Results of cross-validation tests using simulated data under the three species delimitation scenarios presented in Fig. 2. We used varying numbers of SNPs (10, 50 and 1,000; in yellow) and different numbers (10, 50 and 100) and types (blue-Discrete, green-continuous Brownian-Motion (BM), red-continuous Ornstein-Uhlenbeck (OU)) of traits to train the deep learning network architecture presented in Fig. 1. Each bar represents the network accuracy (i.e., probability of choosing the scenario from which the data were simulated) given the input dataset. (a) We present results for each data type separately, (b) detailing the confusion matrices for the data sets with 10 features (in which more classification errors are observed in the lower diagonal, associated with over-lumping) or (c) combining SNPs with one type of trait. For these combined results, we show (d) the weight given to the SNP data branch (the learned weights of the gated concatenation) in the final decision, expressed as a percentage (“% SNPs contribution”) to the final classification decisions.

### Simulated Data: Assessing the Impact of Missing Data

Missing data had a minor effect on classification accuracy (Fig. S1). We observed similar negative effects when introducing random missing values (Fig. S1a and S1b) and when removing samples (Fig. S1c and S1d). Missing data within trait matrices, especially on discrete traits, had a stronger effect when analyzing trait-only data sets than in combined genetic-trait data analyses, where the effect was much smaller (all accuracy values > 95%; Fig. S1a and S1c). In contrast, when we included missing data within the SNP matrices, we observed similar effects for the combined and SNP-only dataset (Fig. S1b and S1d).

### Simulated Data: Speciation Scenarios with Migration

Increasing levels of gene flow (i.e., higher migration rates) among species in the simulated data resulted in lower accuracy by predicting fewer species than we simulated when the network was trained without migration (Fig. 5a). The negative impact of the model violation introduced by higher gene flow was lower when analyzing trait-only data sets or when analyzing combined data sets, particularly at higher migration rates (Fig. 5a, S2a, S3a). We observed a similar trend, although with a smaller effect, when the networks were trained with higher migration but we analyzed simulated data that did not include higher migration rates (Fig. 5b, S2b, S2b). It is important to note that incorporating higher migration into the training procedure resulted in high accuracy levels (>78%) in the trait-only and all combined data sets (Fig. 5c, S2c, S3c). The SNP-only data sets were the most affected by migration: accuracy was lowest at high migration levels, and even when the network was trained under high migration, predictions remained uncertain (e.g., intermediate probabilities across scenarios at *N*_e_m = 0.5; Fig. 5b, 5c). Notably, the weight assigned to the SNP data branch (i.e., the learned weights from the gated concatenation) decreased relative to that assigned to trait data as migration rates increased when higher migration was incorporated during training (Fig. 5, S2 and S3).

**Figure 5.**
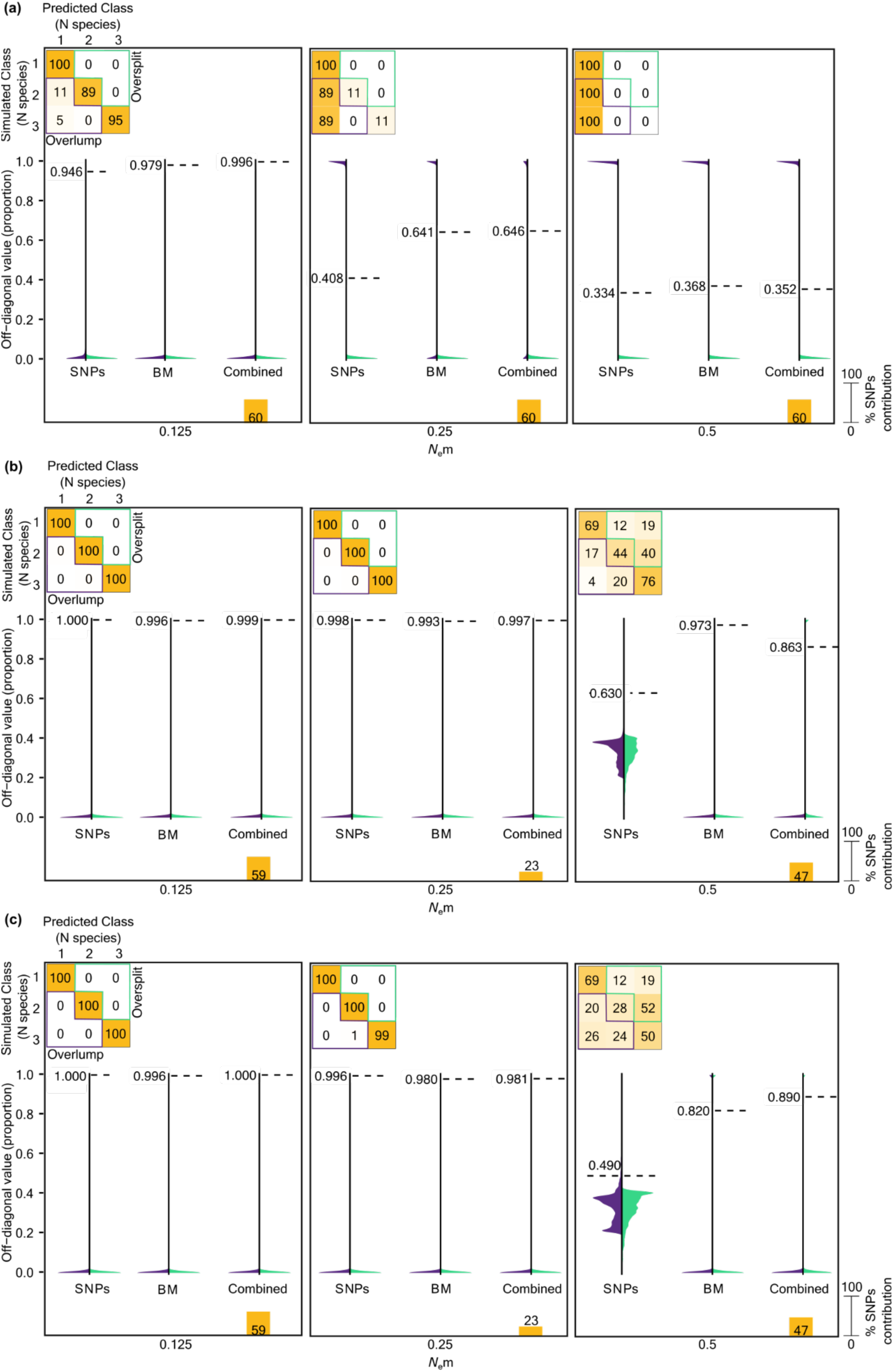
Results of cross-validation tests showing the effect of increasing the level of gene flow, i.e., the migration rate among species pairs, on the accuracy levels of deep learning networks. Results are shown for (a) networks trained with simulated data sets (1,000 SNPs, 100 BM traits) without migration and tested using simulations with migration, for (b) networks trained with migration and tested using simulations without migration, and for (c) networks trained and tested with migration. For each migration rate in each subplot, we show the confusion matrix obtained when comparing the predicted and simulated scenarios for the SNP dataset, showing errors associated with over-lumping in purple at the lower diagonal and errors associated with over-splitting in green at the upper diagonal. We also show a mirrored density plot for each datatype (SNPs, BM traits, and the combined dataset), depicting the distribution of the over-lumping (left wing, in purple) and over-splitting (right wing, in green) errors in predictions. High density at the bottom of the plot represents low error (highly confident correct classifications), while high density at the top represents highly confident incorrect classifications and density in the middle shows ambiguous predictions (intermediate probabilities for all scenarios). The dashed horizontal line indicates the mean classification accuracy (argmax) for each dataset. For the combined results, we also show the weight given to the SNPs data branch (the learned weights of the gated concatenation) in the final decision, expressed as a percentage (“% SNPs contribution”).

### Empirical Data: Sweet Tabaiba

DeepID results for the *Euphorbia balsamifera* species complex indicate that the three-species delimitation scenario proposed by Riina et al. (2021; Fig. 3a.iv and 3a.v) provides the best fit to the data. This result is robust to the type of data used (traits only, SNPs only, or combined data sets) and the evolutionary model assumed for trait evolution (BM or OU), with a higher probability of the scenario including ancient migration (Fig. 3a.v; Table 2). When SNPs were considered alone, the scenario with ancient migration (Fig. 3a.v) had the lower relative probability compared to the other scenarios, with a softmax probability (SP) of 0.905, whereas it showed high probability (SP ∼ 1.0) when traits were considered alone and when SNPs were combined with trait data (Table 2). In all cases, the next best-supported scenario was the one with three species without high migration (Fig. 3a.iv; Table 2). Both models combining SNP and trait data (under BM and OU) showed a higher contribution of SNPs to the gated-concatenation weights (73 and 77%, respectively), as expected given that only four traits were included.

**Table 2.** Model selection results for the two empirical data sets. Softmax probabilities (SP) are given for each scenario and each dataset applied, considering only traits (under BM and OU models), only SNPs, and combined (SNP+traits) data. +M = Migration included.

|  | BM | OU | SNPs | SNPs+BM | SNPs+OU |
| --- | --- | --- | --- | --- | --- |
| Sweet tabaiba |  |  |  |  |  |
| i) One species | 0.000 | 0.000 | 0.000 | 0.000 | 0.000 |
| ii) Two species ((S,B),A) | 0.000 | 0.000 | 0.000 | 0.000 | 0.000 |
| iii) Two species (S,(B,A)) | 0.000 | 0.000 | 0.000 | 0.000 | 0.000 |
| iv) Three species | 0.000 | 0.003 | 0.095 | 0.000 | 0.000 |
| v) Three species+M | 1.000 | 0.997 | 0.905 | 1.000 | 1.000 |
| % SNPs contribution |  |  |  | 73 | 77 |
| Longear Sunfish |  |  |  |  |  |
| i) Two species | 0.000 | 0.000 | 0.000 | 0.000 | 0.000 |
| ii) Six species | 1.000 | 1.000 | 1.000 | 1.000 | 1.000 |
| iii) Six species+M_hybrids | 0.000 | 0.000 | 0.000 | 0.000 | 0.000 |
| iv) Six species+M | 0.000 | 0.000 | 0.000 | 0.000 | 0.000 |
| % SNPs contribution |  |  |  | 42 | 41 |

Cross-validation showed high accuracy for all data sets (Fig. S4), except for distinguishing between the two three-species scenarios (without and with ancient migration, Fig. 3a.iv and 3a.v), which exhibited high levels of confusion in all data sets. This was expected, as these two scenarios are very similar and we only included ancient migration (i.e., occurring from the time of divergence to halfway through the total divergence time). If we disregard the confusion between those two scenarios, the accuracy was very high, especially for SNP-based analyses, with a proportion of simulations assigned for the correctly simulated scenario > 0.99 in both SNP-only and combined data sets (Fig. S4). Accuracy was slightly lower for trait-only analyses, although the simulated scenarios were consistently recovered with SP > 0.84 (Fig. S4) after excluding the two scenarios with three species (Fig. 3a.iv and 3a.v).

### Empirical Data: Longear Sunfish

In the *L. megalotis* species complex, DeepID identified the most likely species delimitation hypothesis as the six-species scenario without high migration, which includes only the baseline interspecific migration (0 < *N*_e_m < 0.1), present in all scenarios (Fig. 3b.ii). This scenario was recovered with high probability (SP = 1.0) across all data sets, regardless of whether only SNPs were considered, SNPs were combined with traits, or only traits were considered. Both combined networks (with BM and OU traits) showed a lower contribution of SNPs to the learned weights of the gated concatenation (42 and 41%, respectively; Table 2) compared to the traits contribution. This was unexpected given the relative size of the data sets (1,000 SNPs versus 31 traits), but is consistent with the results from the simulated data sets when high levels of migration are included during training (Fig. 5c, S2c and S3c).

Cross-validation again showed higher accuracy for the combined data sets (> 70% across all scenarios; Fig. S4), with greater confusion between the two scenarios modeled under higher migration rates (Fig. 3b.iii, 3b.iv). Networks trained using only trait data showed similar accuracy to the combined data sets, whereas the SNP-only network achieved low confusion when distinguishing among scenarios with different numbers of species, but higher confusion among the six-species scenarios (Fig. S4).

## DISCUSSION

Our results show the importance of integrating complementary information from different data types for species delimitation in a flexible framework. The DeepID approach that we developed can integrate data from large genetic data sets with continuous and discrete traits. DeepID is flexible, enabling the incorporation of varying levels of missing values for genetic and trait data. It is also capable of discriminating among complex delimitation hypotheses because it is able to explicitly consider the demographic events associated with the speciation process.

### Simulating the speciation process

An important feature of DeepID is that the inferences are based on coalescent simulations that incorporate important components of the speciation process by including population-level demographic events. Therefore, DeepID coincides with other delimitation methods that incorporate the speciation process through coalescent simulations of genomic data (Camargo et al. 2012; Smith and Carstens 2020; Perez et al. 2022). The main novelty of our approach is the inclusion of explicit thresholds for intraspecific and interspecific divergence times based on the *P*_a_ and *P*_b_ heuristic indexes (Fig. 2) proposed by Rannala and Yang (2020). By taking this approach, DeepID is able to reduce the splitting tendency of coalescent-based methods that frequently delimit intraspecific genetic structure rather than species (Sukumaran and Knowles 2017; Leaché et al. 2019). Cross-validation results for the simple delimitation scenarios (Fig. 4a), the high-migration scenarios (Fig. 5), and the two empirical species complexes analyzed here (Fig. S4) suggest that DeepID is a promising approach for reducing the tendency toward over-splitting commonly observed in MSC-based demographic models. In cases of misclassification, errors more frequently correspond to over-lumping (i.e., predicting fewer species than simulated; lower diagonal of the confusion matrices) than to over-splitting (i.e., predicting more species than simulated; upper diagonal of the confusion matrices). Another strategy recently applied to reduce over-splitting in species delimitation is to use a reference-based framework, in which information from well-studied, related groups or a subset of population lineages is used to establish species limits (Sukumaran et al. 2021; Leaché et al. 2021; Derkarabetian et al. 2022). We outline that the flexibility of our approach makes it possible to easily modify the simulation procedure to use a reference-based strategy when knowledge is available for a closely related group. For example, we could use information from a closely related group to provide priors on the parameters used to derive the custom intra- and interspecific thresholds for *P*_a_ and *P*_b_ in DeepID. In future work, the flexibility of DeepID could be further explored by simulating more realistic migration scenarios. These include episodic migration, such as the scenario recovered for the *E. balsamifera* species complex (Fig. 3a.v; i.e., migration occurring from the initial split of two lineages from a common ancestor to halfway through the total divergence time); scenarios with decreasing migration, in which gene flow is strongest at divergence and gradually or exponentially reduces towards the present; and temporally explicit, piecewise models in which migration is governed by the opening and closing of dispersal barriers. Additional realism could be achieved by incorporating heterogeneous selection across loci; unaccounted selection may lead to false inferences of gene flow in demographic analyses by increasing heterogeneity in levels of divergence or mimicking the effect of changes in population size (Smith and Hahn 2024). Conversely, the use of unphased data (sequence contigs) in our analysis of the *E. balsamifera* species complex may have led to a conservative estimation of genetic diversity. By collapsing diploid genotypes into consensus sequences, rare variants may be filtered out, leading to lower estimates of effective population size (*N*_e_) and migration rates. Despite this, DeepID recovered a scenario with gene flow (episodic migration), indicating that migration signals remain detectable even when using unphased contigs.

### Using deep learning to integrate data and compare species delimitation hypotheses

DeepID integrates genomic and trait data simulated under competing species delimitation scenarios. By using a CNN to perform pattern recognition and discover meaningful features from an “image” of the genomic data (Torada et al. 2019; DeGiorgio et al. 2026), we avoid the limitations of other simulation-based methods that rely on summarizing genetic information through a set of user-specified statistics (e.g., ABC or MLP) and thus may fail to capture important features of the data (Perez et al. 2022; Perez and Gascuel 2025). Moreover, combining the detected features from genomic data with those extracted from the trait data into a unified framework results in a flexible approach that can accommodate different levels of missing data (Fig. S1) and is more robust to model misspecification than genomic data alone (Fig. 5). Species delimitation studies, especially those that integrate different data sources, often face the problem of incomplete data, as information is not always available for every single sample and all datatypes. Notably, DeepID recovered the correct scenario with high accuracy even when large proportions of missing values were introduced, both as randomly scattered entries across the matrix (Fig. S1a and S1b) and as a complete absence of data for subsets of samples (Fig. S1c and S1d). Combining complementary information from different data types (SNPs and continuous and discrete traits), which may reflect different stages of the speciation process (Carstens et al. 2013), can improve the realism and accuracy of species delimitation hypotheses (Solís-Lemus et al. 2015; Eberle et al. 2019; Pyron et al. 2026). In most comparisons, the combined approach showed accuracy similar to or higher than that obtained using genomic or trait data alone (Fig. 4), especially when model assumptions were violated by incorporating higher migration rates among species (Fig. 5). Accordingly, an important result from DeepID is that under model violation (i.e., with the inclusion of high migration), both trait-only and combined SNP+trait models achieved higher accuracy than models based solely on genomic data (Fig. 5). We attribute this to the relative robustness of trait data to coalescent noise (e.g., incomplete lineage sorting), which affects more the SNP data under high migration or large *N*_e_ scenarios but has a less pronounced effect on traits as a result of the approach used to model them (i.e., trait evolution under a BM, OU, or Mk model provides a signal that is partially independent of gene tree discordance).

### Reassessing species delimitation hypotheses for empirical systems

To test the predictive performance and potential limitations of DeepID, we used both simulated and empirical examples of species delimitation hypotheses. For the latter, we selected two case studies (one plant and one animal species complex) representing varying levels of complexity in their speciation scenarios and previously studied by specialists. These studies provide an expected species delimitation hypothesis, i.e., an expert-based species identification that has been translated (or could be translated) into a new formal taxonomy (Riina et al. 2021; Kim et al. 2022). These serve as a form of ground truth (Borowiec et al. 2022), allowing us to assess whether DeepID can recover the same speciation scenarios identified by experts (Riina et al. 2021; Kim et al. 2022).

DeepID supported, with a high probability (Table 2), the species delimitation hypotheses published in the original studies (i.e., three species for the *E. balsamifera* species complex and six species for the *L. megalotis* species complex) and was flexible enough to accommodate different sample sizes, data types, numbers and types of traits, and varying levels (and patterns) of missing data. The *E. balsamifera* plant species complex represents a simple speciation scenario. Riina et al. (2021), based on a combination of phylogenomic data (Villaverde et al. 2018), morphometric characters associated with leaf shape and size (traditionally used by *Euphorbia* taxonomists; Molero et al. 2002), and climatic niche analysis, recognized the existence of three species with deep phylogenetic splits. *E.sepium* diverged from its sister clade (*E. adenensis-E. balsamifera*) circa 6.9 million years ago (Ma), whereas the latter two species separated around 3.5 Ma; no evidence of secondary gene flow among the three species has been detected in population phylogenomic analyses (Villaverde et al. 2018; Rincón-Barrado et al. 2024). This pattern of sequential divergence among three currently reproductively isolated species (Fig. 3a.iv and 3a.v) was also recovered by DeepID, with the highest support for the scenario including higher levels of ancient migration restricted to an early divergence interval following species splitting (Fig. 3av). This scenario was consistently recovered with high probability across all data sets analyzed (trait-only, SNP-only, and combined; Table 2).

The *Lepomis megalotis* species complex represents a less straightforward speciation setting. Previous results from morphological analyses coupled with multispecies coalescent species delimitation, demographic modeling, and admixture assessment (Kim et al. 2022) suggested the presence of six species with secondary gene flow among several species pairs after the initial divergence. Again, our approach was able to recover the expected scenario with the divergence of six species (Fig. 3b.ii). Surprisingly, DeepID predictions assigned low support to the two scenarios assuming high post-divergence gene flow (Fig. 3b.iii, 3b.iv), while the low gene flow scenario (six species with only baseline migration, Fig. 3b.ii) received the highest probability across all analyzed data sets (Table 2). Although this stands in contrast with an extensive hybridization speciation hypothesis, we believe that our results suggest that the baseline migration levels (*N*_e_m < 0.1) are enough to account for the “secondary contact or rare hybridization events between nonsister species” (Kim et al. 2022) in the *L. megalotis* species complex, without the need to include higher migration levels (0.1 < *N*_e_m < 0.5).

Convolutional neural networks are increasingly being employed in automated species identification due to their capacity to use feature maps and visual image recognition to classify samples into discrete classes (Borowiec et al. 2022; Karbstein et al. 2024). However, CNN use in systematics for defining species boundaries, and especially for species delimitation based on demographic modeling of speciation scenarios, is less frequent (Perez et al. 2022). Here, we demonstrate that CNNs can be used to compare alternative speciation delimitation hypotheses and to assess how robust the predictions are to violations of model assumptions, such as interspecific gene flow after the initial divergence (Smith and Carstens 2020; Kim et al. 2022). Also, our results provide support for the idea that incorporating phenotypic data (e.g., discrete and continuous traits) along with genomic data sets can help in species delimitation, especially for scenarios involving speciation with gene flow or those that fall in the gray zone of species delimitation (de Queiroz 2007), as taxa might diverge along other evolutionary dimensions (morphology, ecology) before achieving genetic reciprocal monophyly (Solís-Lemus et al. 2015; Sukumaran et al. 2021).

## CONCLUSIONS

We developed an integrative deep learning approach (DeepID) that combines genomic and trait information via convolutional neural networks and multilayer perceptrons into a unified artificial neural network framework for species delimitation and the comparison of different speciation scenarios. The predictive performance (accuracy) of our deep neural network was higher for data sets combining genomic and trait data than when we analyzed each data type alone. Therefore, our results support previous suggestions that using multiple sources of data (e.g., continuous and discrete traits, together with genetic information) accounts for different stages of the speciation process and can help to reduce the over-splitting bias observed in popular multispecies coalescent methods. Furthermore, our method is flexible, allowing us to accommodate the inherent complexity of empirical species delimitation data sets that use multiple sources of information with varying levels and configurations of missing data.

## DATA AVAILABILITY STATEMENT

The two empirical data sets used in this study, as well as all scripts and notebooks, are available from https://github.com/manolofperez/DeepID and Dryad (http://datadryad.org/share/AS3W9OC-L7HqV4FZ7HbeG_JzZNEGRqcvcRec_rCGQlc). A tutorial on how to apply DeepID to empirical systems is available on https://github.com/manolofperez/DeepID/blob/master/EmpiricalData/Tutorial_DeepID/Tutorial_DeepID.ipynb.

## ACKNOWLEDGEMENTS

We would like to appreciate the valuable feedback of the EvoGenomics.ai network (https://www.evogenomics.ai/) members to improve this work. We also acknowledge the HPC infrastructure at the Federal University of Sao Carlos (Cluster UFSCar).

