## Supplementary material for "Integrative taxonomy using traits and genomic data for Species Delimitation with Deep learning": Supp. Figure

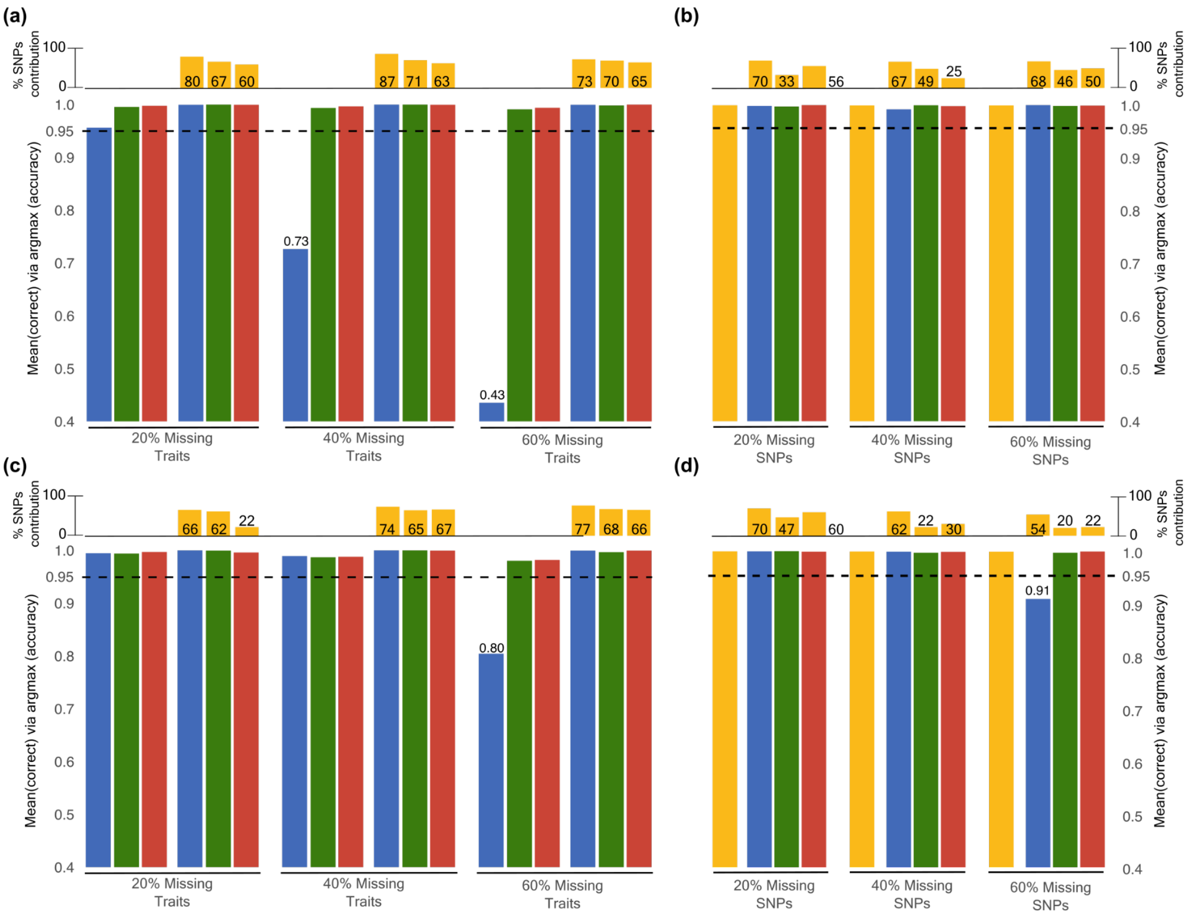


**Figure S1.** Results of cross-validation tests showing the effect of increasing the percentage of missing data on the accuracy levels of deep learning networks trained with simulated datasets (1,000 SNPs and 100 traits) under the three species delimitation scenarios presented in Fig. 2. We randomly added a proportion (20, 40 and 60% of missing values (0 entries) to the (a) trait and (b) SNP data matrices or replaced all entries in these same proportion of samples with missing values for (c) traits and (d) SNPs. Each bar represents the network accuracy (i.e., probability of choosing the scenario from which the data were simulated) given the input dataset. We present results for each data type separately (blue-Discrete, green-continuous Brownian-Motion (BM), red-continuous Ornstein-Uhlenbeck (OU), yellow- SNPs). The results of networks trained with combined data sets of SNPs and traits are depicted with a yellow bar above, which represents the weight given to the SNP data branch (the learned weights of the gated concatenation) in the final decision, expressed as a percentage ("% SNPs contribution") to the final classification decisions.

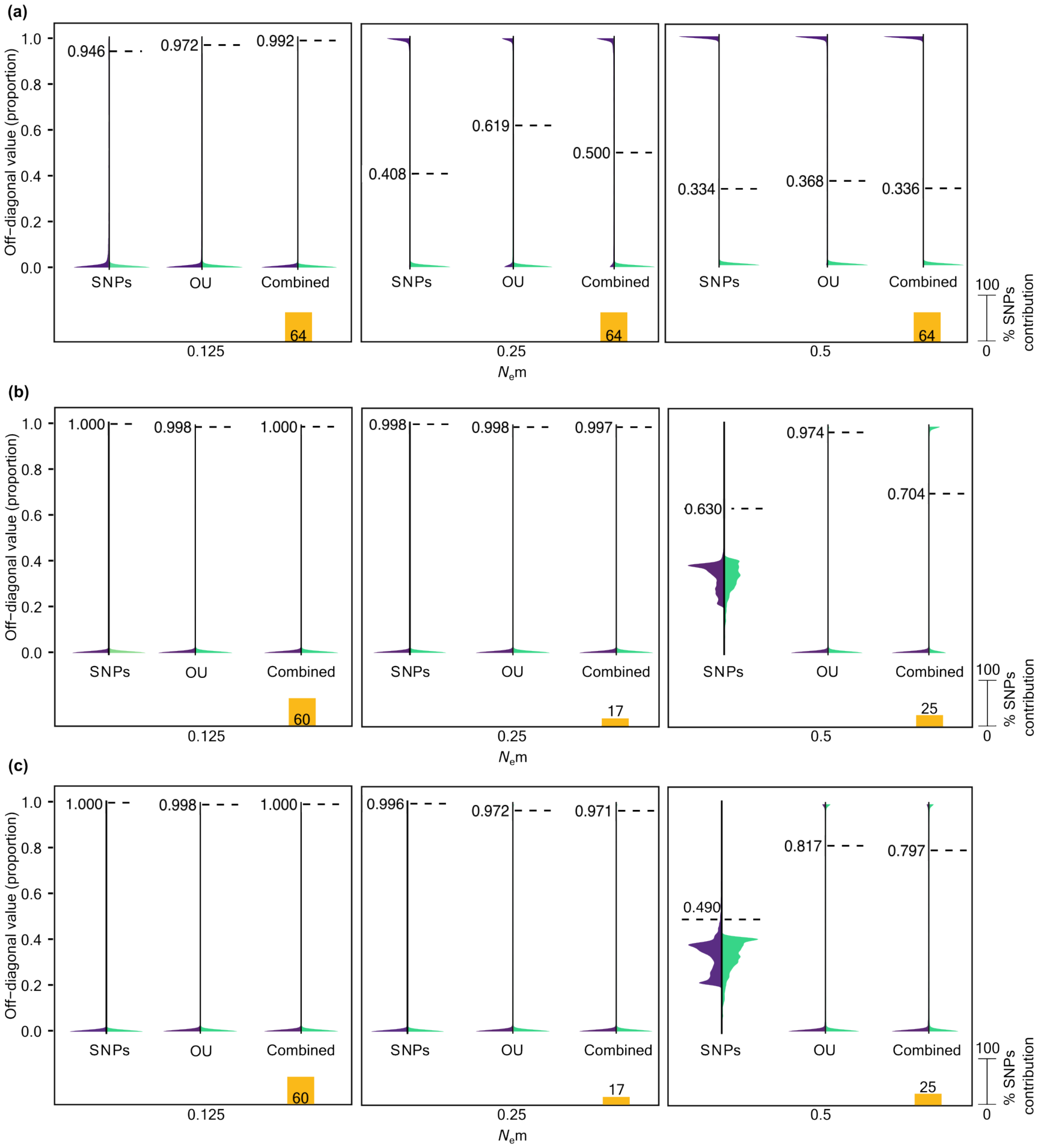


**Figure S2.** Results of cross-validation tests showing the effect of increasing the level of gene flow, i.e., the migration rate among species pairs, on the accuracy levels of deep learning networks. Results are shown for (a) networks trained with simulated data sets (1,000 SNPs, 100 OU traits) without migration and tested using simulations with migration, for (b) networks trained with migration and tested using simulations without migration, and for (c) networks trained and tested with migration. For each migration rate in each subplot, we show a mirrored density plot for each datatype (SNPs, OU traits, and the combined dataset), depicting the distribution of the over-lumping (left wing, in purple) and over-splitting (right wing, in green) errors in predictions. High density at the bottom of the plot represents low error (highly confident correct classifications), while high density at the top represents highly confident incorrect classifications and density in the middle shows ambiguous predictions (intermediate probabilities for all scenarios). The dashed horizontal line indicates the mean classification accuracy (argmax) for each dataset. For the combined results, we also show the weight given to the SNPs data branch (the learned weights of the gated concatenation) in the final decision, expressed as a percentage (“% SNPs contribution”).

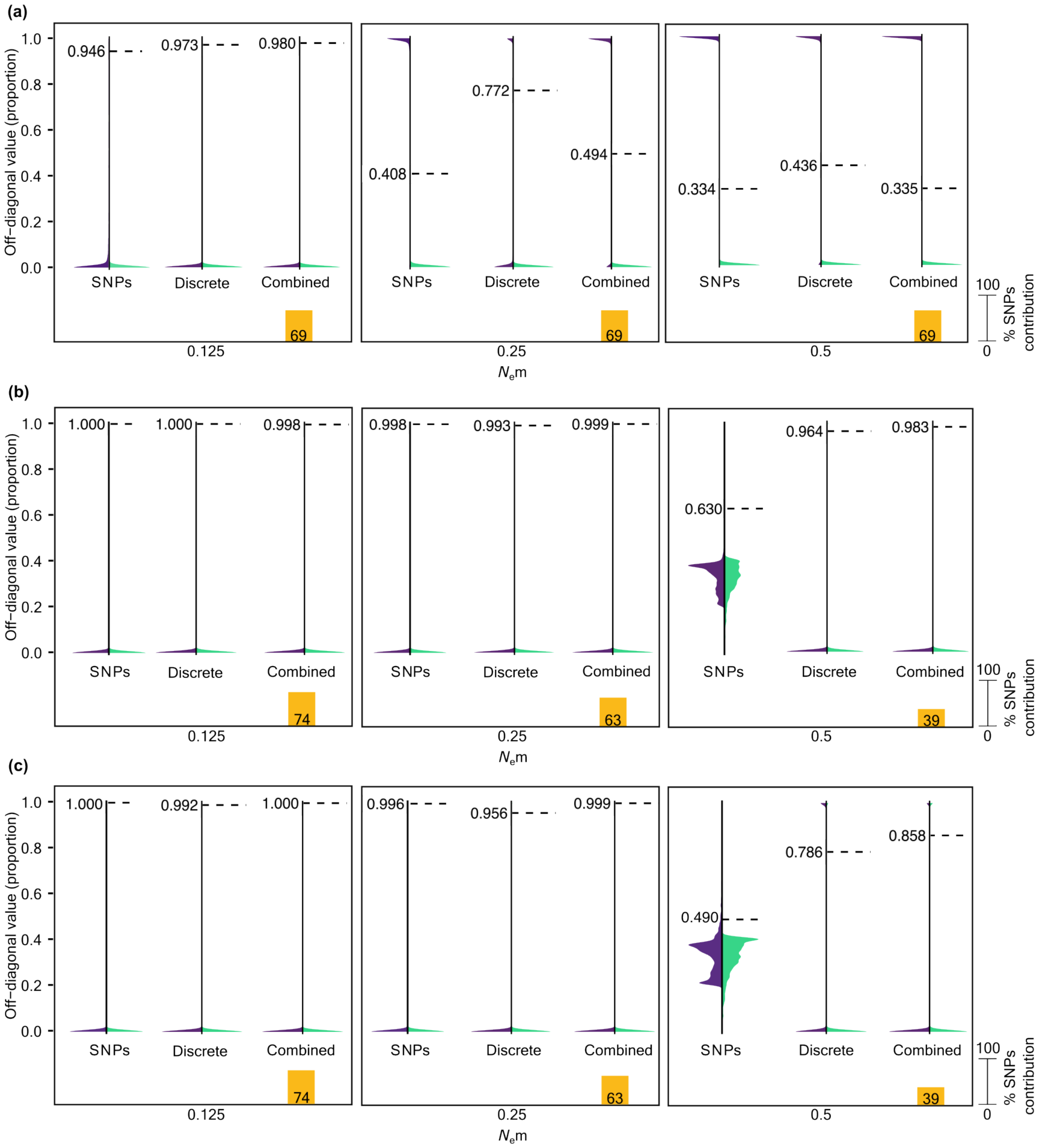


**Figure S3.** Results of cross-validation tests showing the effect of increasing the level of gene flow, i.e., the migration rate among species pairs, on the accuracy levels of deep learning networks. Results are shown for (a) networks trained with simulated data sets (1,000 SNPs, 100 discrete traits) without migration and tested using simulations with migration, for (b) networks trained with migration and tested using simulations without migration, and for (c) networks trained and tested with migration. For each migration rate in each subplot, we show a mirrored density plot for each datatype (SNPs, discrete traits, and the combined dataset), depicting the distribution of the over-lumping (left wing, in purple) and over-splitting (right wing, in green) errors in predictions. High density at the bottom of the plot represents low error (highly confident correct classifications), while high density at the top represents highly confident incorrect classifications and density in the middle shows ambiguous predictions (intermediate probabilities for all scenarios). The dashed horizontal line indicates the mean classification accuracy (argmax) for each dataset. For the combined results, we also show the weight given to the SNPs data branch (the learned weights of the gated concatenation) in the final decision, expressed as a percentage ("% SNPs contribution").

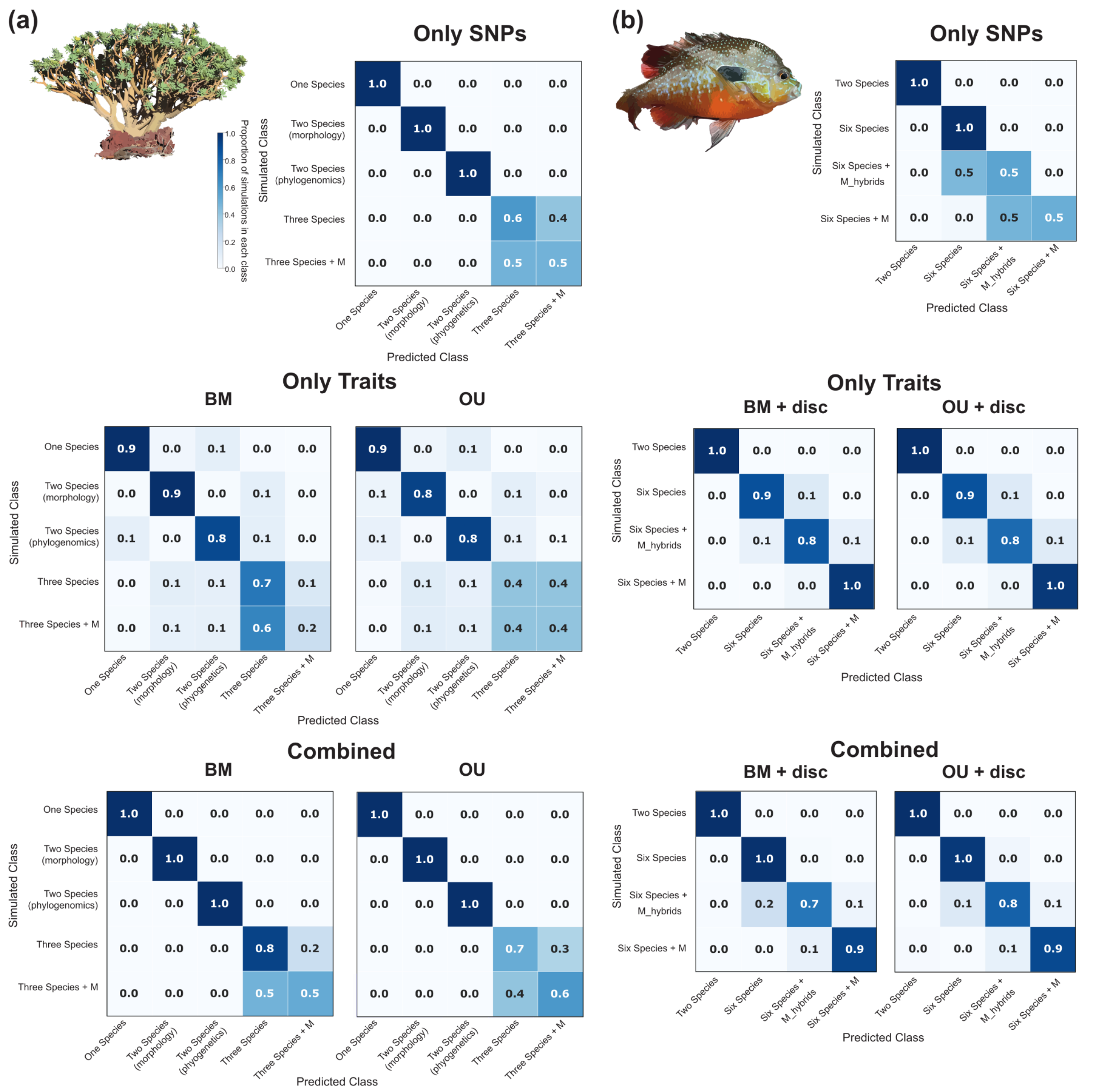


**Figure S4.** Confusion matrices showing the percentage of predicted classes for each simulated scenario in (a) the sweet tabaiba (*Euphorbia balsamifera* species complex) and (b) the longear sunfish (*Lepomis megalotis* species complex) data sets using test data (independent simulations not evaluated in the training step). Species were illustrated by one of the authors (MFP) in Adobe Illustrator CC 2018.
